# Challenges to case-only analysis for gene-environment interaction detection using polygenic risk scores: model assumptions and biases in large biobanks

**DOI:** 10.1101/2025.09.15.676316

**Authors:** Wenmin Zhang, Qiongshi Lu, Tianyuan Lu

**Author notes:** Correspondence to Wenmin Zhang and Tianyuan Lu.

## Abstract

Gene-environment interaction is important for studying complex diseases. Case-only analysis has been proposed to improve power for GxE detection. However, case-only analysis relies on key assumptions, including correct specification of the disease risk model and marginal independence between genetic and environmental variables. In this study, we systematically investigate the challenges of case-only analysis using polygenic risk scores (PRS) as genetic variables in large biobanks. Through simulations, we demonstrate that the false positive control of PRS-based case-only analysis depends on the log-linear disease risk model and weak main effects, and that it is prone to false positives under other commonly used disease risk models. We then conduct case-only analyses for breast cancer, prostate cancer, class 3 obesity, and short stature in the UK Biobank, using PRS derived from non-overlapping chromosome sets (e.g., even-numbered and odd-numbered chromosomes) that are unlikely to interact with each other. The resulting case-only regression estimates consistently show negative shifts compared to population-based estimates, suggesting false positives driven by collider bias due to model misspecification. Furthermore, correlations between chromosome set-specific PRS, likely driven by assortative mating or population stratification, suggest additional sources of confounding. Our results underscore the challenges of applying PRS-based case-only analysis in large biobank settings and highlight the need for caution when interpreting case-only results.

## 1 Introduction

Gene-environment interactions (GxE) is important for understanding the multifactorial nature of complex diseases [1]. However, classical regression-based approaches for GxE analysis often have limited statistical power. Several statistical methods have been developed to improve power for GxE detection [2, 3, 4, 5, 6, 7]. One of the earliest and most widely discussed approaches is case-only analysis [8, 9, 10, 11, 12]. Specifically, under the assumption that a genetic variable and an environmental variable are independent in the study population, their association among cases may be used to detect the multiplicative GxE effect on the risk of a disease [8, 9, 10, 11, 12, 13].

With the establishment of large biobanks and the availability of genome-wide association study (GWAS) summary statistics, case-only analysis has been extended to genome-wide settings by using polygenic risk scores (PRS) as the genetic variable [14]. Assuming a log-linear model for disease risk, PRS-based case-only analysis estimates the interaction between a PRS and an environmental variable by regressing the PRS on the environmental variable among cases [14]. This approach benefits from improved power, as a PRS can aggregate information across many genetic variants.

However, the increased power comes with caveats. Importantly, deviation from the log-linear disease risk model may lead to spurious association between PRS and environmental variable among cases due to collider bias, even in the absence of a true interaction effect (**Figure** 1A). For instance, the logistic model quantifies disease risk on the log-odds scale and is routinely used in GWAS for binary disease outcomes[15, 16, 17, 18], while the liability threshold model assumes disease manifests only when a latent liability exceeds a threshold [19]. Since these commonly used models do not conform to the log-linear model, it remains unclear whether PRS-based case-only analysis yields valid results across the a broad range of complex traits and diseases.

**Figure 1:**
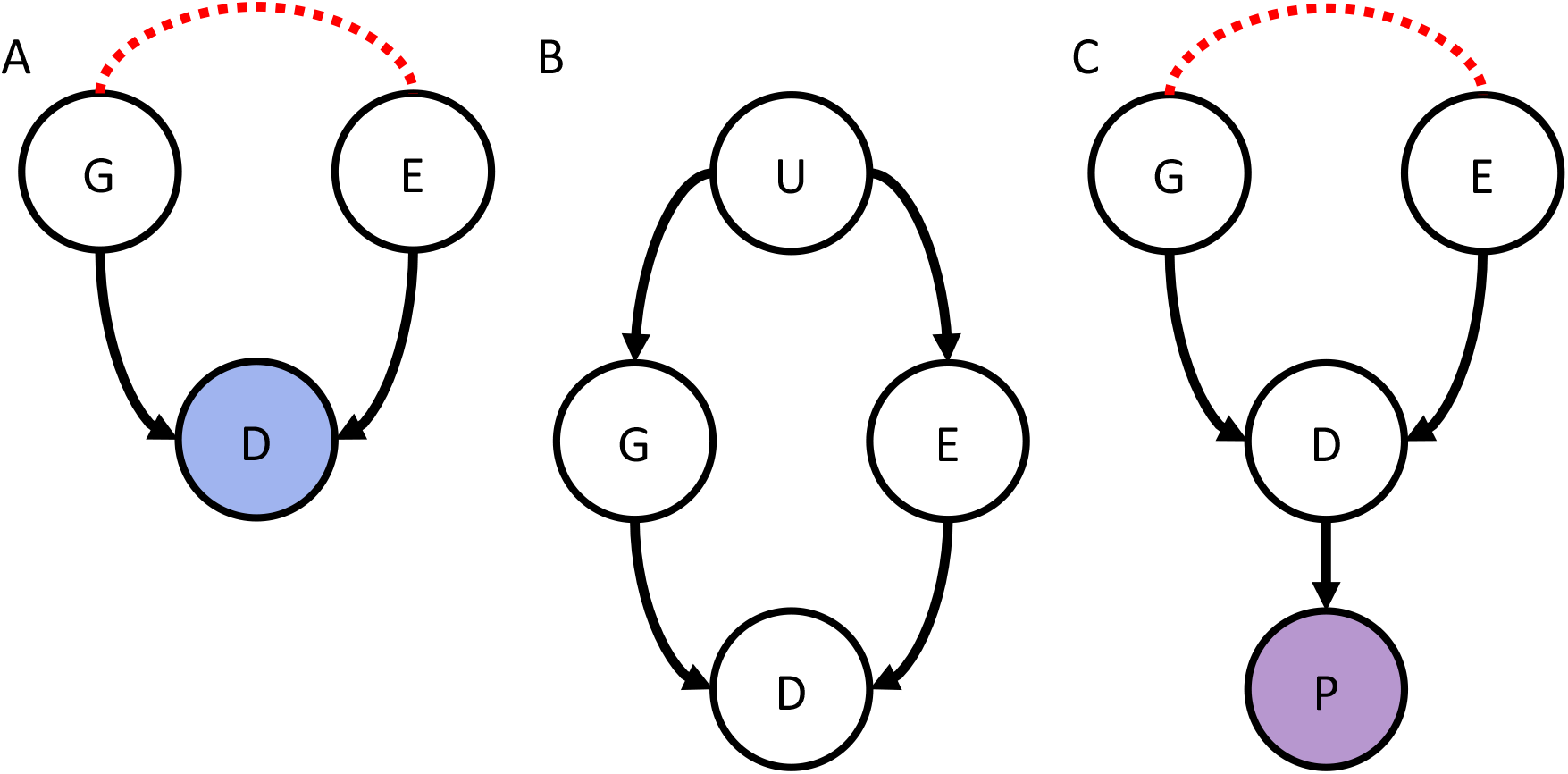
Challenges of PRS-based case-only analysis in large biobanks. **A**: Conditioning on disease status (*D*) creates a collider structure between PRS (*G*) and environmental variable (*E*), which may lead to false positives in case-only analysis when the log-linear model for disease risk does not hold. **B**: Confounders (*U*) can induce a spurious association between PRS (*G*) and environmental variable (*E*), violating the marginal independence assumption in case-only analysis. **C**: Participation (*P*) bias associated with disease status (*D*) can also create a collider structure that leads to spurious associations between PRS (*G*) and environmental variable (*E*), violating the marginal independence assumption in case-only analysis. Colored nodes represent variables being conditioned upon. Red dotted lines indicate conditional associations. Importantly, these challenges are not unique to PRS-based studies and may generalize to all case-only designs relying on similar assumptions.

Furthermore, the fundamental assumption of marginal independence between the PRS and the environmental variable in the study population may not hold in large biobanks. Many environmental variables, including lifestyle and socioeconomic factors, are themselves partially heritable, which can induce correlation between the PRS and the environmental variables, thereby violating the independence assumption. Although methods to adjust for the heritability of environmental factors have been proposed, such as including an additional PRS for the environmental factor in the case-only analysis [14], residual confounding may still persist (**Figure** 1B). Sources of such confounding include complex population structure and assortative mating [20]. Additionally, disease status may influence an individual’s likelihood of participating in biobank studies, potentially creating a collider structure and leading to spurious associations, even if the PRS and environmental variable are marginally independent in the general population (**Figure** 1C).

In this study, we systematically investigate the challenges of PRS-based case-only analysis in large biobanks. We conduct simulations to assess the performance of case-only analysis under three common disease risk models: piecewise log-linear, logistic, and liability threshold models. This allows us to quantify how violations of log-linear assumption can lead to false positives. To empirically test for spurious associations, we implement a chromosome-partitioning strategy in the UK Biobank [21], in which we construct PRS from non-overlapping chromosome sets (e.g., even-numbered vs. odd-numbered chromosomes) that are unlikely to have strong biological interactions. We treat one PRS as the genetic variable and the other as a proxy for an environmental variable to evaluate whether they become spuriously correlated in case-only analysis. This analysis is performed across four disease outcomes: breast cancer, prostate cancer, class 3 obesity, and short stature. Our results underscore the importance of carefully evaluating the assumptions underlying PRS-based case-only analysis and highlight the challenges of applying this approach to large-scale biobank data.

## 2 Methods

### 2.1 False positives in PRS-based case-only analysis due to collider bias

We demonstrate that collider bias will lead to false positives in PRS-based case-only analysis, except in the special case of a log-linear model for disease risk. Under the null hypothesis of no interaction effect, we assume a general disease risk model:

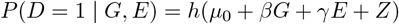

where *D* is the binary disease status, *G* is the PRS, *E* is the environmental variable, *µ*_0_ is a baseline parameter, *β* is the effect size of PRS, *γ* is the effect size for environmental variable, and *Z* captures unobserved factors affecting disease risk.

In the log-linear model, we have:

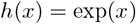

In the logistic model, we have:

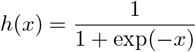

In the liability threshold model, we have:

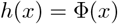

where Φ(*x*) is the cumulative distribution function of the standard normal distribution.

In the absence of interaction effect, the conditional distribution of PRS (*G*) among cases (*D* = 1) should be independent of the environmental variable (*E*). Here, we show that this property only holds under the log-linear disease risk model. We have the conditional distribution of *G* among cases (*D* = 1) as follows:

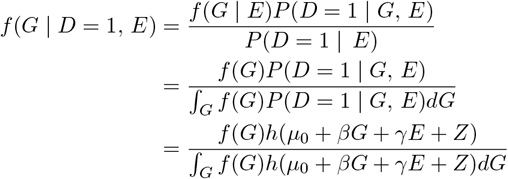

In order for *f* (*G* | *D* = 1, *E*) to be independent of *E, h*(*µ*_0_ + *βG* + *γE* + *Z*) must be multiplicatively separable in *G* and *E*, i.e., it can be expressed as a product of a function of *G* and a function of *E*:

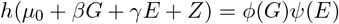

such that *ψ*(*E*) in the denominator cancels out with *ψ*(*E*) in the numerator.

This requirement leads to a functional equation known as the Cauchy’s exponential equation, which, under mild regularity conditions, has the exponential function as its only viable solution [22]. Thus, the log-linear model is the unique case (within this class) where the independence holds.

As an illustration, under the log-linear model and assuming 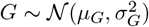, the distribution of *G* among cases simplifies to:

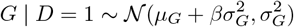

which is indeed independent of *E*.

However, for other disease risk models, such as the logistic and liability threshold models, the conditional distribution of *G* does depend on *E*, which can lead to false positives in PRS-based case-only analysis. We demonstrate this by illustrating conditional probability density functions *f* (*G* | *D* = 1, *E*) under the logistic model and liability threshold model in **Figure** 2.

**Figure 2:**
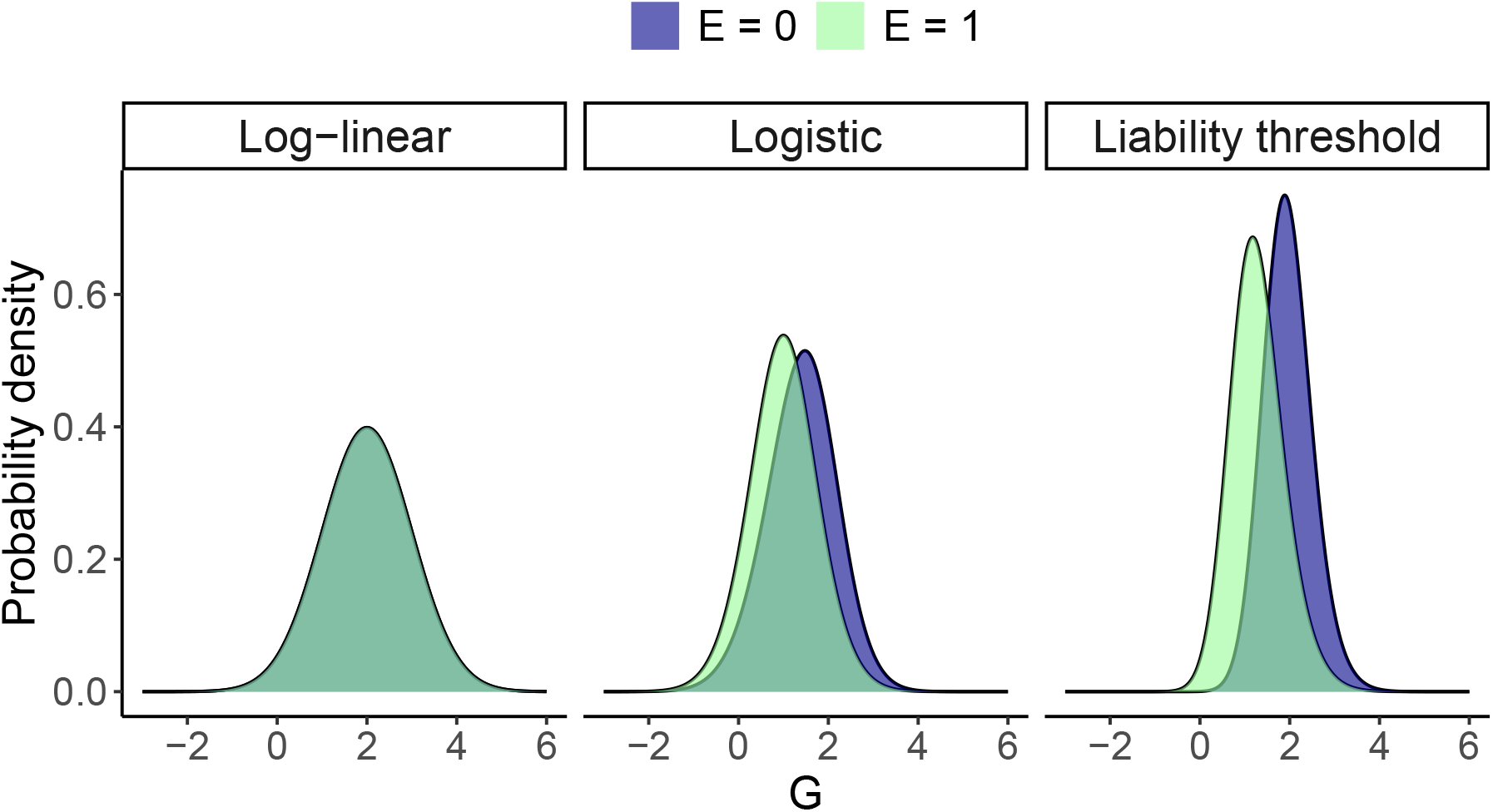
Conditional distributions of PRS among cases under different disease risk models. This is a simulated example without a true interaction effect, where *G* ~ 𝒩 (0, 1) and *p*(*D* = 1 | *G, E*) = *h*(−4+2*G*+2*E*). Under the log-linear model, the conditional distribution of PRS (*G*) among cases (*D* = 1) is independent of the environmental variable (*E*). However, under the logistic model and liability threshold model, the conditional distribution of PRS (*G*) among cases (*D* = 1) depends on the environmental variable *E*.

### 2.2 Simulation study

We next conducted simulations to quantify the false positive rate in case-only analysis under different disease risk models without a true interaction effect. For each model, we generated synthetic datasets with sample sizes of *N* = 100,000 and *N* = 500,000 to investigate whether larger sample sizes exacerbate the false positive rate.

PRS (*G*), environmental variable (*E*) and the additional disease risk-influencing variable (*Z*) were independently sampled from standard normal distributions with zero mean and unit variance. For simplicity, we set the genetic and environmental effect sizes to be equal, i.e., *β* = *γ*. The disease status was then sampled according to probabilities under the log-linear, logistic, and liability threshold models, respectively.

For the log-linear model, to ensure disease probabilities remained in the valid range [0, 1], we used a bounded version of the exponential function:

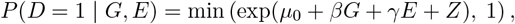

To ensure comparability across models, we numerically selected model-specific baseline parameter *µ*_0_ and effect sizes so that:

- Disease prevalence was approximately 1% across all simulation settings and across all models;
- The marginal associations of *G* with *D* and *E* with *D* were approximately equal across all models, as measured by coefficients from univariate logistic regressions of *D* on *G* and *D* on *E*.

We evaluated a range of effect sizes, where the approximate marginal logistic regression coefficients of *D* on *G* and *D* on *E* varied from 0.1 to 1.0 in increments of 0.1. These model parameters are summarized in **Supplementary Table** S1.

Each combination of data-generating model, effect size, and sample size was replicated 10,000 times. Within each simulation replicate, we regressed *G* on *E* among cases (*D* = 1) using linear regression. The empirical false positive rate was computed as the proportion of simulations where the p-value was below 0.05 or 0.005. We also examined quantilequantile plots of p-values, where an inflation factor (λ) *>* 1.1 was considered indicative of substantial false positive inflation.

### 2.3 UK Biobank

The UK Biobank is a large, population-based cohort of approximately 500,000 individuals aged 40-69 years at recruitment between 2006 and 2010 [21]. Participants were identified through National Health Service records and invited to attend one of the 22 assessment centers across the United Kingdom, where they completed questionnaires on demographics, lifestyle, and medical history, underwent physical assessments, and provided biological samples. Health outcomes were ascertained through linkage to hospital inpatient and death records. Baseline anthropometric measurements, including height and body mass index (BMI), were collected using standardized protocols by trained staff.

Genotyping was conducted using the UK BiLEVE Axiom and UK Biobank Axiom arrays, followed by centralized quality control [21]. Genotypes were imputed using the Haplotype Reference Consortium panel [23], with additional coverage from the UK10K and 1000 Genomes Phase 3 reference panels [24, 25].

For this analysis, we included participants of genetically inferred European ancestry based on principal component analysis (data field 22006). To minimize bias from related individuals, we randomly excluded one individual from each pair with a kinship coefficient ≥0.0442, as estimated using KING [26]. This threshold corresponds approximately to third-degree relatives.

### 2.4 Case definitions

Breast cancer cases were defined based on any of the following: an International Classification of Diseases (ICD)-9 code of 174 or an ICD-10 code of C50 in hospital inpatient or death records, or a self-reported history of breast cancer (data field 20001, code 1002). Only female participants were considered for this outcome.

Prostate cancer cases were defined based on any of the following: an ICD-9 code of 185 or an ICD-10 code of C61 in hospital inpatient or death records, or a self-reported history of prostate cancer (data field 20001, code 1044). Only male participants were considered for this outcome.

Class 3 obesity was defined as a BMI of 40 kg/m^2^ or higher, measured at the baseline visit. This threshold corresponds to the World Health Organization’s definition of class 3 obesity, which is associated with significantly elevated risks of cardiometabolic and other health complications [27, 28].

Short stature was defined using baseline height measurements. To account for age-related variation, height was first regressed on age separately in males and females. The resulting residuals were standardized within each sex to produce z-scores with zero mean and unit variance [29]. Short stature was defined as a z-score below −2.25 [30], consistent with the US Food and Drug Administration’s definition, which corresponds approximately to the lowest 1.2% of the population distribution.

### 2.5 Polygenic risk scores

We calculated PRS for breast cancer, prostate cancer, BMI, and height based on large-scale GWAS that did not include UK Biobank participants, to avoid overfitting due to sample overlap.

For breast cancer and prostate cancer, we used publicly available PRS from the PGS Catalog [31]: PGS000335 [32] for breast cancer and PGS002240 [33] for prostate cancer. These scores were derived from GWAS of individuals of European ancestry, including 76,192 cases and 63,082 controls for breast cancer [34], and 79,148 cases and 61,106 controls for prostate cancer [35]. Both scores were developed using the PRS-CS method [36]. The European ancestry population from the 1000 Genomes Phase 3 data [25] was used as the linkage disequilibrium (LD) reference panel.

For BMI and height, we developed PRS using GWAS summary statistics from the GIANT consortium [37, 38], restricted to European ancestry individuals (N = 322,154 for BMI and 253,288 for height). We applied a clumping and thresholding approach with a p-value threshold of 1 *×* 10^−5^ and an LD *r*^2^ threshold of 0.01, using the European ancestry population from the 1000 Genomes Phase 3 data [25] as the LD reference panel. The weights for the selected variants included in the PRS are provided in **Supplementary Tables** S2 and S3.

Per-chromosome PRS were calculated using imputed genotype dosages in UK Biobank participants with PLINK v1.9 [39].

### 2.6 Statistical analyses

To illustrate the potential challenges in case-only analysis, we partitioned the genome into two non-overlapping sets of chromosomes: one containing all odd-numbered autosomes and the other containing all even-numbered autosomes, effectively treating the former as the genetic variable and the latter as the environmental variable. For each individual, we computed set-specific PRS by aggregating the per-chromosome scores within each chromosome set.

We first verified that each set-specific PRS was significantly associated with the corresponding disease outcome using logistic regression, adjusting for age at recruitment, assessment center, genotyping array, the first 10 genetic principal components, and sex (only for class 3 obesity and short stature).

Next, we assessed the association between the odd-numbered chromosome set score and the even-numbered chromosome set score separately in the full study population and the subset of cases only using linear regression. We conducted these association tests both with and without adjustment for the same covariates above to investigate the impact of population stratification.

To contextualize these observed associations and generate empirical distributions, we extended this analysis by randomly assigning 11 autosomes to form set 1 and the remaining 11 to form set 2. We then regressed PRS derived from set 1 (treated as the genetic variable) on PRS derived from set 2 (treated as the environmental variable), repeating the association tests in both the full study population and among cases only, with and without covariate adjustment, across 5,000 random partitions. This approach allows for evaluation of the expected variability in association estimates. All scores were standardized to have zero mean and unit variance prior to analysis. Under the assumption of independent assortment, two chromosome set-specific PRS should be independent in the full study population. A detectable association in the full study population may indicate systematic bias, for example, due to assortative mating [40].

Furthermore, in the absence of true inter-chromosomal interactions, the distribution of association estimates in case-only analysis should be approximately centered at zero. If there are true inter-chromosomal interactions influencing disease risk, the ability to detect them in the case-only analysis depends on whether the interacting chromosomes are assigned to the same or different sets. Specifically, when interacting chromosomes are within the same set, the interaction effect may not be detected in the set–set association. If interacting chromosomes are split across different sets, the interaction may induce a detectable association between the set-specific scores among cases. As a result, repeated random partitioning of chromosomes could lead to bimodal or multimodal distributions of association estimates in the case-only analysis. In contrast, a unimodal distribution of association estimates that consistently deviates from zero in the case-only analysis may suggest systematic bias.

## 3 Results

### 3.1 Simulation results

Simulation results demonstrated that case-only analysis was more likely to maintain appropriate false positive control when the underlying model was approximately log-linear (**Figure** 3). In contrast, in the logistic or liability threshold models, false positive rates became inflated, especially as effect sizes increased (**Figure** 3).

**Figure 3:**
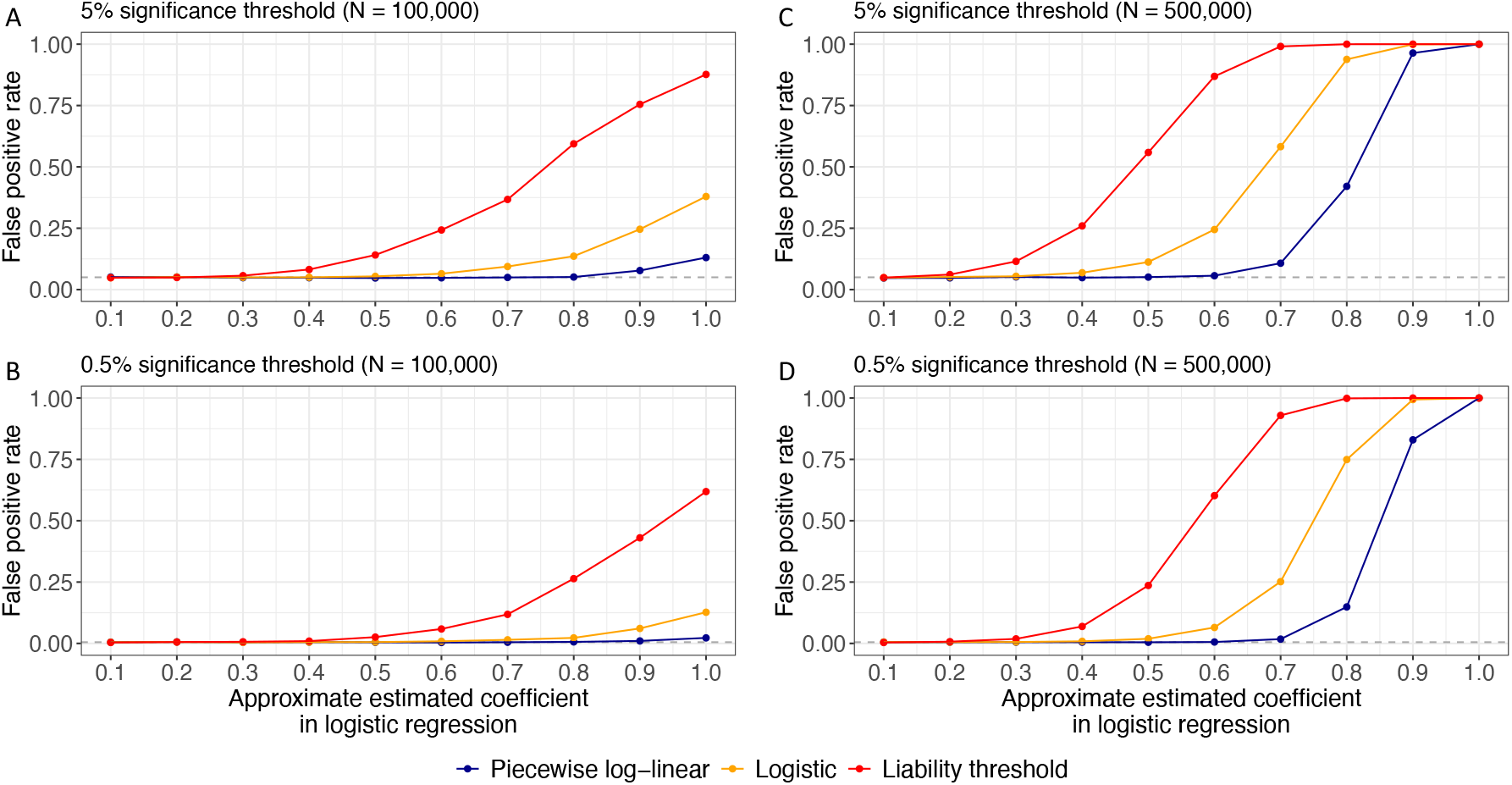
False positive rates in case-only analysis in simulations without true interaction effect. False positive rates are shown at **A**: 5% significance level, sample size = 100,000; **B**: 0.5% significance level, sample size = 100,000; **C**: 5% significance level, sample size = 500,000; **D**: 0.5% significance level, sample size = 500,000. False positive rates are evaluated under three disease risk models: piecewise log-linear, logistic, and liability threshold. The x-axis indicates the effect sizes of the polygenic risk score (*G*) and environmental variable (*E*), measured by their estimated coefficients from logistic regression. The y-axis represents the observed false positive rate in simulations. Grey dashed horizontal lines mark nominal significance thresholds.

Specifically, with a sample size of 100,000, false positive control was adequate for effect sizes ≤ 0.3 across all models (**Figure** 3A and B and **Supplementary Figure** S1). However, inflation of false positive rate became evident at larger effect sizes: ≥ 0.4 for the liability threshold model, ≥ 0.6 for the logistic model, and ≥ 0.9 for the piecewise log-linear model. Increasing the sample size to 500,000 further exacerbated this issue (**Figure** 3C and D and **Supplementary Figure** S2), with inflation of false positive rate emerging at effect sizes as low as 0.2 for liability threshold model, 0.4 for logistic model, and 0.7 for piecewise log-linear model.

### 3.2 PRS for breast cancer, prostate cancer, BMI, and height in UK Biobank

The UK Biobank study population consisted of 335,993 unrelated individuals of European ancestry, including 180,767 females and 155,226 males (**Supplementary Table** S4). The average age was 56.7 years (SD = 7.9) for females and 57.1 years (SD = 8.1) for males. Breast cancer was present in 8.3% of females, while prostate cancer was present in 7.2% of males. Mean baseline BMI was 27.0 kg/m^2^ (SD = 5.1) for females and 27.8 kg/m^2^ (SD = 4.2) for males, with class 3 obesity identified in 2.3% and 1.3% of females and males, respectively. Mean height was 162.7 cm (SD = 6.2) in females and 176.0 cm (SD = 6.8) in males, with short stature identified in approximately 1.1% of females and 1.2% of males. Missing data were minimal across all variables (≤0.3%).

We confirmed the associations between the PRS and their corresponding outcomes in the UK Biobank (**Figure** 4A and **Supplementary Tables** S5 and S6). For breast cancer, a one standard deviation increase in the whole-genome PRS was associated with an odds ratio (OR) of 1.77 (95% CI: 1.74–1.80). When the PRS was constructed using only odd-numbered chromosomes, the association was attenuated but remained substantial (OR = 1.42; 95% CI: 1.40–1.45), and similarly for even-numbered chromosomes (OR = 1.55; 95% CI: 1.52–1.57). A similar pattern was observed for prostate cancer, where a one standard deviation increase in the whole-genome PRS was associated with an OR of 2.11 (95% CI: 2.07–2.16), while the odd- and even-numbered chromosome set-specific PRS had ORs of 1.65 (95% CI: 1.62-1.68) and 1.70 (95% CI: 1.66-1.73), respectively (**Figure** 4A; **Supplementary Table** S5).

**Figure 4:**
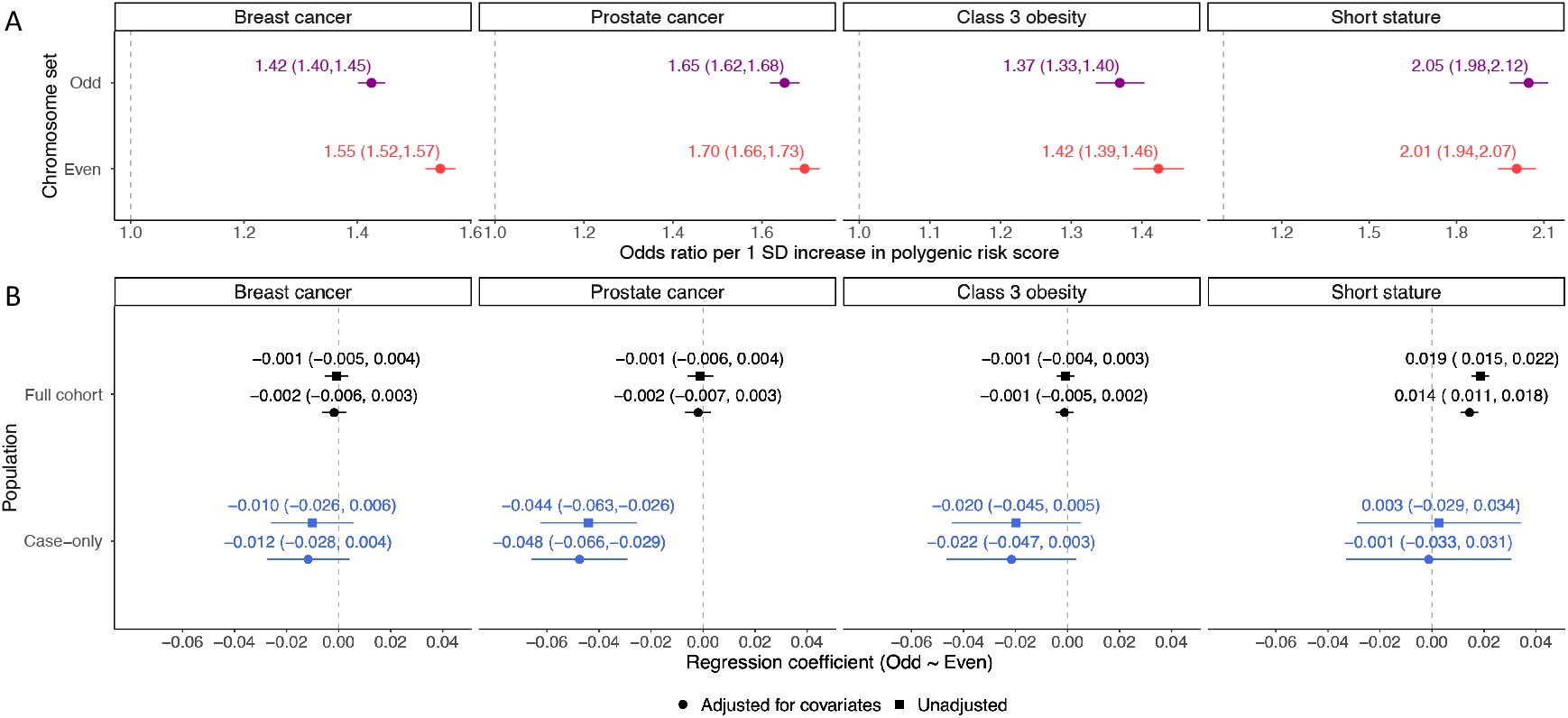
Association tests using PRS derived from odd and even chromosomes. **A**: Odds ratios for disease outcomes associated with PRS derived from odd- and even-numbered chromosomes. **B**: Linear regression coefficients for the association between PRS derived from odd-numbered chromosomes and PRS from even-numbered chromosomes, estimated in the full study population and in cases only. Error bars represent 95% confidence intervals. Vertical dashed lines denote the null value.

For anthropometric traits, the BMI PRS explained 2.6% of the variance in BMI in the full study population, with similar contributions from the odd-numbered chromosomes (*R*^2^ = 1.2%) and even-numbered chromosomes (*R*^2^ = 1.4%; **Supplementary Table** S6). As expected, a one standard deviation increase in the whole-genome BMI PRS was associated with an increased odds of class 3 obesity (OR = 1.61; 95% CI: 1.57–1.65), with similar effects from the odd-numbered chromosomes (OR = 1.37; 95% CI: 1.33–1.40) and even-numbered chromosomes (OR = 1.42; 95% CI: 1.39–1.46). The height PRS showed a strong association with height (*R*^2^ = 8.6%), again with similar contributions from odd-numbered chromosomes (*R*^2^ = 4.4%) and even-numbered chromosomes (*R*^2^ = 4.3%). A one standard deviation decrease in the whole-genome height PRS was associated with an increased odds of short stature (OR = 2.73; 95% CI: 2.64–2.82), with similar effects from the odd-numbered chromosomes (OR = 2.05; 95% CI: 1.98–2.12) and even-numbered chromosomes (OR = 2.01; 95% CI: 1.94–2.07; **Figure** 4A and **Supplementary Table** S6).

### 3.3 Associations between odd- and even-numberd chromosome set-specific PRS

We assessed the association between PRS derived from odd- and even-numbered chromosomes (**Figure** 4B). In the full study population, the unadjusted regression coefficients were −0.001 for breast cancer (95% CI: −0.005, 0.004), −0.001 for prostate cancer (95% CI: −0.006, 0.004), and −0.001 for class 3 obesity (95% CI: −0.004, 0.003). After adjusting for age at recruitment, assessment center, genotyping array, the first 10 genetic principal components, and sex (for class 3 obesity), the regression coefficients were marginally more negative: −0.002 for breast cancer (95% CI: −0.006, 0.003), −0.002 for prostate cancer (95% CI: −0.007, 0.003), and −0.001 for class 3 obesity (95% CI: −0.005, 0.002; **Figure** 4B).

Importantly, in the case-only analysis, the regression coefficients were substantially more negative, although not necessarily statistically significant. Specifically, the unadjusted regression coefficients were −0.010 for breast cancer (95% CI: −0.026, 0.006), −0.044 for prostate cancer (95% CI: −0.063, −0.026), and −0.020 for class 3 obesity (95% CI: −0.045, 0.005). The regression coefficients after adjustment for the same covariates were −0.012 for breast cancer (95% CI: −0.028, 0.004), −0.048 for prostate cancer (95% CI: −0.066, −0.029), and −0.022 for class 3 obesity (95% CI: −0.047, 0.003; **Figure** 4B).

Height PRS showed a different pattern. In the full study population, there was a substantial positive association between the odd- and even-numbered chromosome set–specific PRS. The unadjusted regression coefficient was 0.019 (95% CI: 0.015, 0.022), which marginally attenuated after covariate adjustment (0.014; 95% CI: 0.011, 0.018). In contrast, among individuals with short stature, the unadjusted regression coefficient was 0.003 (95% CI: −0.029, 0.034), and the adjusted estimate was −0.001 (95% CI: −0.033, 0.031; **Figure** 4B).

### 3.4 Distrbution of inter-chromosomal polygenic risk score association estimates

To further assess the association of chromosome set–specific PRS, we examined the distributions of regression coefficients based on 5,000 random partitions of the autosomes into two sets. All resulting distributions of regression coefficients appeared to be unimodal (**Figure** 5). Detailed regression coefficients for each random partition are provided in **Supplementary Table** S7.

**Figure 5:**
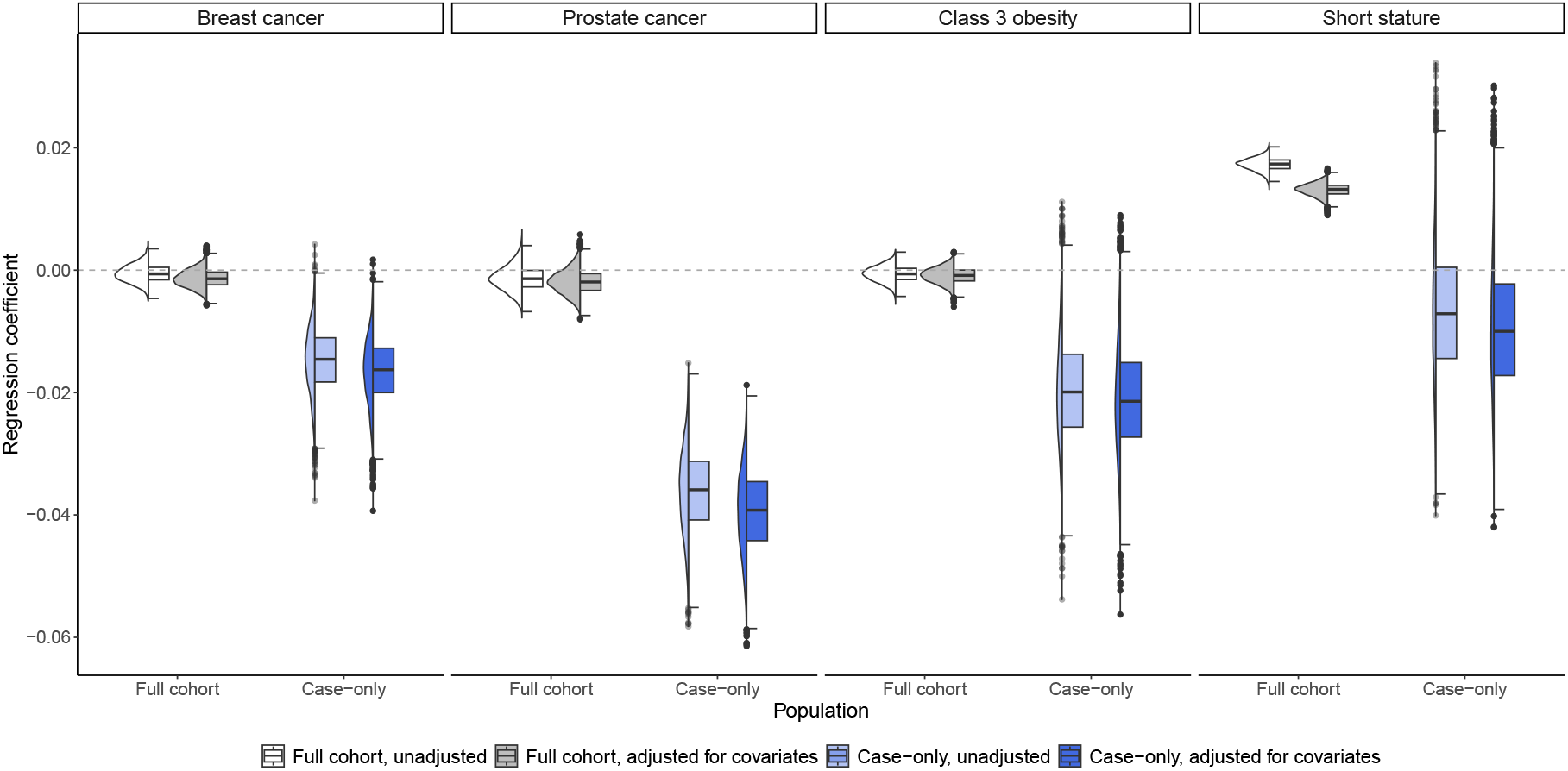
Empirical distributions of association estimates between PRS derived from two-set partitions. Violin plots and boxplots display the distributions of association estimates between PRS derived from two-set partitions of the autosomes, based on 5,000 random replicates. The distributions are shown separately for the full study population and for cases, with and without adjustment for covariates. In each boxplot, the central box represents the interquartile range (IQR), the horizontal line inside the box indicates the median, whiskers extend to 1.5 times the IQR or to the most extreme data points within this range, and individual points beyond the whiskers are shown as outliers. The violin plots illustrate the kernel density estimate of the distributions.

In the full study population, regression coefficients were centered near zero for breast cancer, prostate cancer, and class 3 obesity, consistent with the minimal correlations observed in the odd- and even-numbered chromosome set analyses. For breast cancer, the regression coefficients had a median of −0.001 (inter-quartile range, IQR −0.005 to 0.004) without adjustment and −0.002 (IQR −0.006 to 0.003) after covariate adjustment. For prostate cancer, the median was −0.001 (IQR −0.006 to 0.004) without adjustment and −0.002 (IQR −0.007 to 0.003) with covariate adjustment. For class 3 obesity, the median was −0.001 (IQR −0.004 to 0.003) without adjustment and −0.001 (IQR −0.005 to 0.002) after adjustment (**Figure** 5).

In contrast, the case-only regression coefficient distributions shifted substantially toward negative values. For breast cancer, the regression coefficients had a median of −0.015 (IQR −0.026 to −0.006) without adjustment and −0.017 (IQR −0.028 to −0.007) after covariate adjustment. For prostate cancer, the median was −0.036 (IQR −0.063 to −0.026) without adjustment and −0.039 (IQR −0.066 to −0.029) with adjustment. For class 3 obesity, the median was −0.020 (IQR −0.045 to −0.014) without adjustment and −0.021 (IQR −0.047 to −0.015) after adjustment (**Figure** 5).

Notably, for short stature, in the full study population, the regression coefficients had a median of 0.017 (IQR 0.015 to 0.022) without adjustment and 0.013 (IQR 0.011 to 0.018) after covariate adjustment. Among individuals with short stature, the median was −0.007 (IQR −0.029 to 0.034) without adjustment and −0.001 (IQR −0.033 to 0.031) with adjustment (**Figure** 5).

## 4 Discussion

Studying GxE effects is important for understanding complex diseases and inform targeted strategies for disease prevention and treatment [1]. Among available methods, the case-only analysis offers appealing statistical efficiency, but it depends critically on key assumptions, including a log-linear disease risk model, which differs from the routinely adopted logistic model in GWAS for binary disease outcomes. In this work, through both simulations and real data analyses in the UK Biobank, we demonstrated potential issues in using PRS as the genetic variable for case-only analysis in large biobanks.

In simulations, case-only analysis performed best under the piecewise log-linear model, as expected. In contrast, false positive rates were inflated under both the logistic and liability threshold models, with the most severe inflation observed under the liability threshold model. Across all models, the extent of inflation increased with larger main effect sizes and larger sample sizes. These results highlight that the validity of case-only analysis depends strongly on correctly specifying the disease risk model, which cannot be validated in real data analyses. Therefore, there is a trade-off between power and false positive control in case-only analysis. Specifically, when main effects are small, false positive rates tend to be well-controlled. However, interaction effects are generally unlikely to be sufficiently strong to be reliably detected in such scenarios, because empirically, strong interaction effects without strong main effects are rare [41, 42, 43]. Conversely, when main effects are larger and interaction effects may be more detectable, false positive rates are more likely to be inflated, especially under model misspecification. The practical relevance of this trade-off requires further consideration, particularly in the context of complex traits with uncertain genetic architectures.

We then applied a chromosome-partitioning strategy to empirically evaluate the performance of PRS-based case-only analysis in the UK Biobank. Although we treated one chromosome set as the environmental variable in this analysis, the issues identified here are likely generalizable to actual environmental variables. This approach has the advantage that, in the absence of other sources of bias, PRS derived from non-overlapping sets of chromosomes should be marginally independent. Additionally, there is currently no strong evidence of inter-chromosomal interaction effects for the diseases analyzed in this study. Consistent with this, the distributions of regression coefficients across 5,000 random chromosome partitions appeared unimodal. As a result, in the case-only analysis, regression coefficients for chromosome set–specific PRS showed substantial negative shifts compared to those from the full study population across all chromosome partitions and diseases. These results are consistent with simulation findings under the logistic and liability threshold models. The liability threshold model, in particular, may better describe the underlying data-generating processes for class 3 obesity and short stature, since they were defined by dichotomizing continuous traits. The observed pattern suggests that collider bias could contribute to false positives in case-only analysis.

Additional challenges may arise when working with large biobanks and cohorts such as the UK Biobank. First, we observed a substantial positive association between chromosome set-specific PRS for height in the full study population. This result is consistent with the known signature of assortative mating [40], whereby individuals with similar heights are more likely to mate, leading to correlated distributions of height-increasing or height-decreasing alleles across chromosomes. Such correlations can result in spurious associations in both population-based and case-only analyses.

Second, the observation that covariate adjustment resulted in more negative regression coefficients in both the full population and case-only analyses suggests that unadjusted analyses may be affected by confounding factors inducing spurious positive correlations between the two chromosome set-specific PRS. Notably, even after adjusting for measurable confounders such as demographic variables, genotyping array, and genetic principal components, subtle population stratification and cryptic relatedness remain difficult to fully eliminate in practice.

Third, although the regression coefficients for breast cancer, prostate cancer, and class 3 obesity were centered near zero in the full study population, they were marginally negative. This pattern might reflect previously recognized participation bias in the UK Biobank [44, 45, 46]. Additionally, survival bias may be present, since individuals must live long enough and remain healthy enough to participate. If the disease status influences the probability of study participation, spurious correlations may emerge.

Taken together, these findings underscore the need for caution in PRS-based case-only analysis. Associations observed in case-only designs may result from various non-biological factors, including collider bias due to model misspecification or data collection processes, as well as confounding specific to the study population. As a result, statistical significance in case-only analysis does not necessarily imply a mechanistic interaction [13]. Drawing valid conclusions requires conducting multiple sensitivity analyses under alternative modeling assumptions and triangulating multiple lines of evidence.

## Acknowledgements

This research has been conducted using the UK Biobank Resource under Application Number 440468. T.L. has been supported by start-up funding from the Office of the Vice Chancellor for Research and Graduate Education, School of Medicine and Public Health, and Department of Population Health Sciences at the University of Wisconsin-Madison. The funders have no role in study design; collection, management, analysis and interpretation of data; or the decision to submit for publication. We thank Dr. Celia M. T. Greenwood for insightful discussions.

## 5 Declarations

### 5.3 Ethics Approval

This research was conducted using secondary data and was deemed exempt by the Institutional Review Board at the University of Wisconsin–Madison. For analyses involving the UK Biobank, ethical approval is covered by the UK Biobank’s designation as a Research Tissue Bank, granted by the North West Multi-centre Research Ethics Committee. This framework allows researchers to conduct analyses under the existing approval without the need for additional project-specific ethical review, except in cases such as participant re-contact.

### 5.4 Author Contributions

T.L. and W.Z. conceptualized the study, curated the data, performed the analyses, and wrote the original draft. All authors interpreted the results and reviewed and edited the paper critically.

### 5.5 Consent to participate

Not applicable.

### 5.6 Consent to publish

Not applicable.

