## Supplementary Figures for "Challenges to case-only analysis for gene-environment interaction detection using polygenic risk scores: model assumptions and biases in large biobanks"

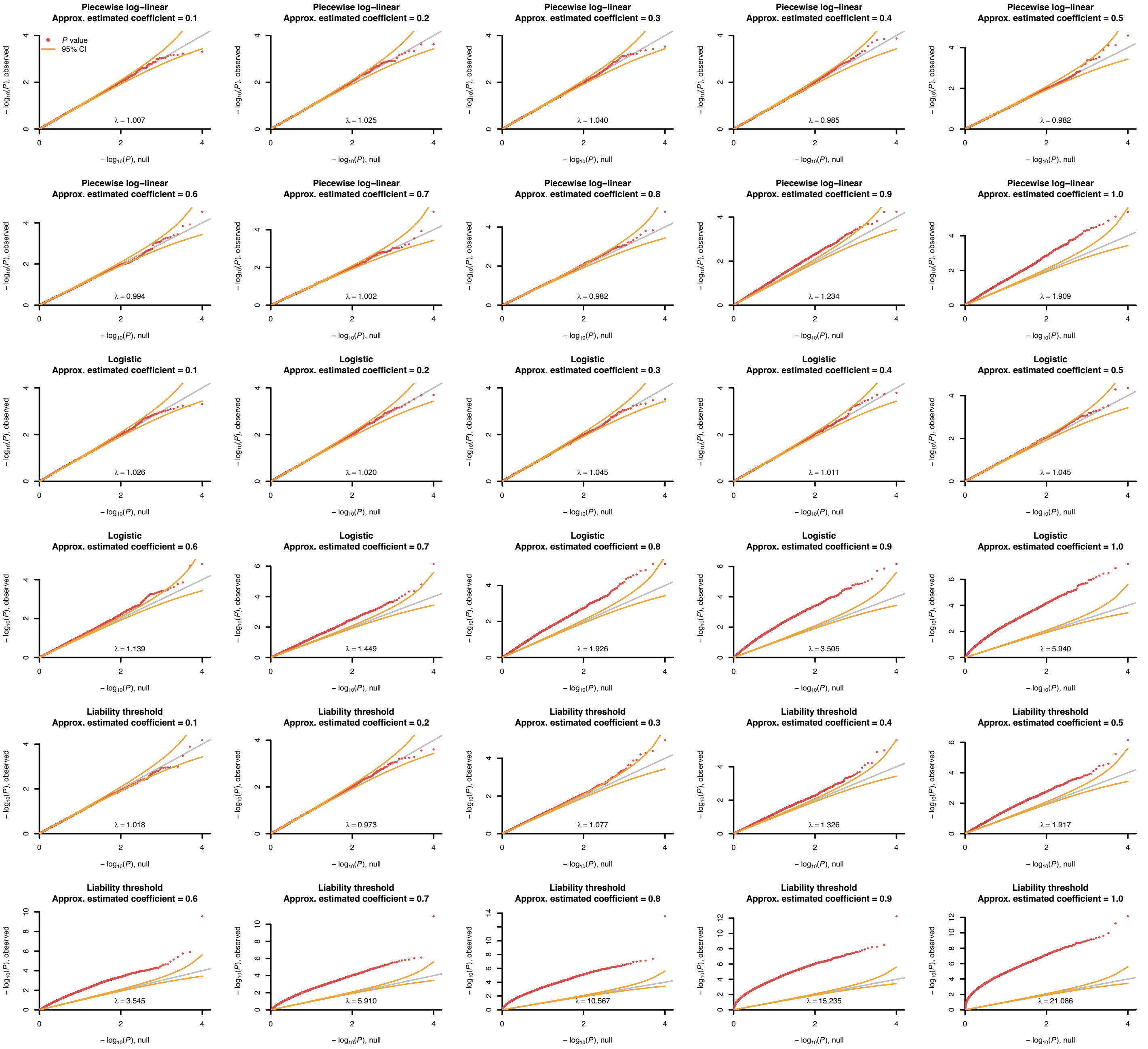


Supplementary Figure S1. Quantile–quantile plots of p-values from case-only analyses (sample size = 100,000) under the null hypothesis. Three models were evaluated: piecewise log-linear, logistic, and liability threshold. Simulations were run for varying effect sizes, each replicated 10,000 times. Red dots show observed p-value distributions. Grey lines indicate a slope of 1. Orange curves denote 95% confidence intervals for well-calibrated p-values. Inflation factors ($\lambda$) are indicated.

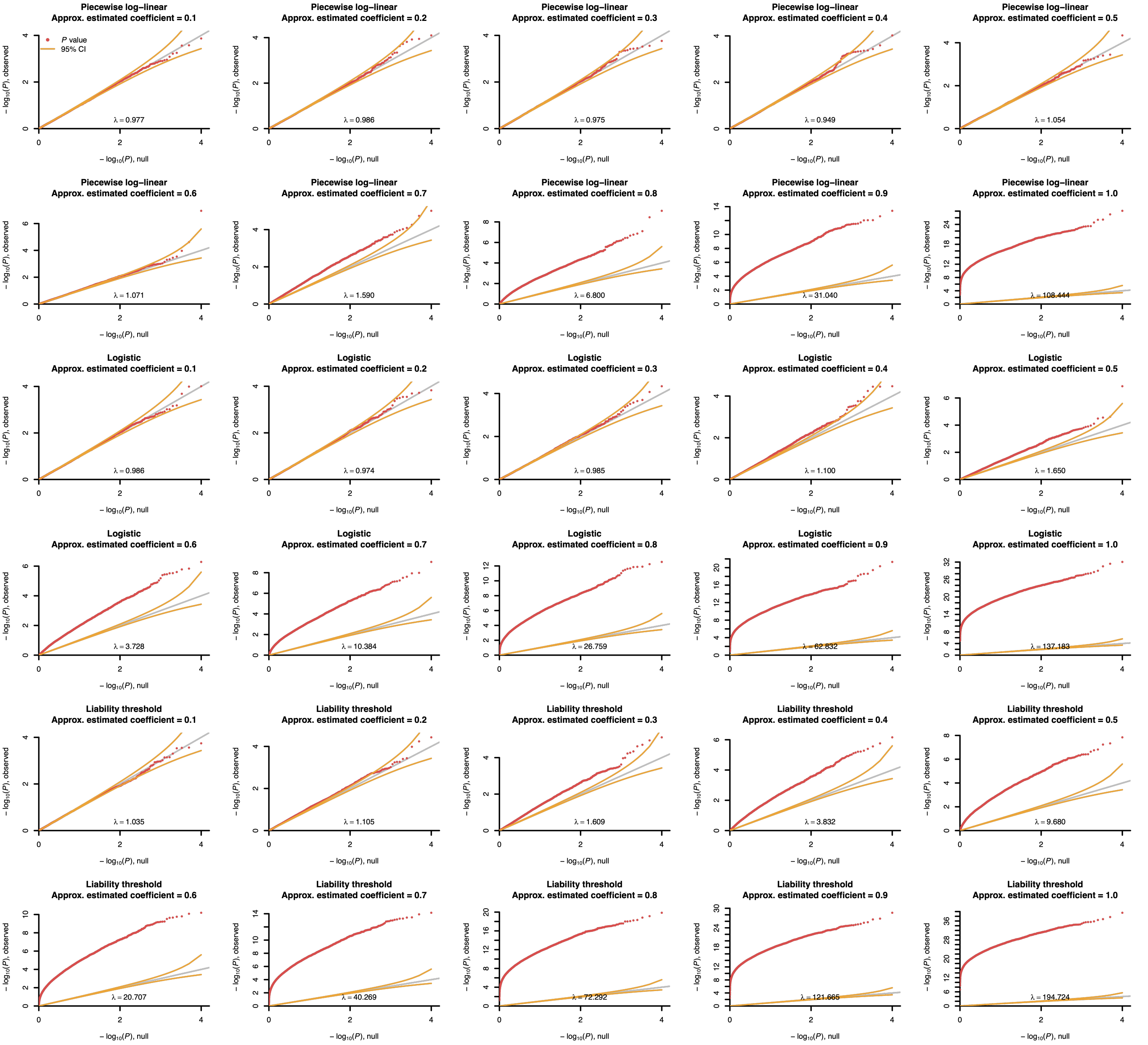


Supplementary Figure S1. Quantile–quantile plots of p-values from case-only analyses (sample size = 500,000) under the null hypothesis. Three models were evaluated: piecewise log-linear, logistic, and liability threshold. Simulations were run for varying effect sizes, each replicated 10,000 times. Red dots show observed p-value distributions. Grey lines indicate a slope of 1. Orange curves denote 95% confidence intervals for well-calibrated p-values. Inflation factors ($\lambda$) are indicated.
